# In silico analysis reveals differential targeting of enterovirus species by commonly used PCR assays

**DOI:** 10.1101/2024.09.13.612945

**Authors:** Van N. Trinh, Nisha Mulakken, Kara L. Nelson, Nicholas Be, Rose S. Kantor

**Affiliations:** Department of Civil and Environmental Engineering, University of California, Berkeley, CA, USA; Computing and Global Security Directorates, Lawrence Livermore National Laboratory, Livermore, CA, USA; Physical and Life Sciences Directorate, Lawrence Livermore National Laboratory, Livermore, CA, USA

**Keywords:** qPCR, assay signature erosion, enterovirus, wastewater-based epidemiology

## Abstract

Quantitative polymerase chain reaction (qPCR) assays are sensitive molecular tools commonly used to quantify pathogens in environmental samples. These assays have become a staple of wastewater-based surveillance and often form the basis of quantitative microbial risk assessments. However, PCR assays may fail to capture all of their intended targets due to signature erosion over time. Here, we performed an *in silico* analysis of four human enterovirus PCR assays to assess signature erosion against the NCBI virus database. The predicted number of genomes hit by each assay rose alongside total genomes in the database through 2010 but the proportion of predicted hits began to level off with the emergence of the clinically important enterovirus D-68. We found that although all assays captured a majority of enterovirus species, only one recently developed assay adequately captured enterovirus D species. Some assays also appeared more likely to capture non-human enteroviruses than others, an important consideration for data interpretation. We conclude that broad-spectrum virus assays applied to environmental samples should be regularly checked against expanding sequence databases and provide methods to do so.

## Introduction

Quantitative and digital polymerase chain reaction assays (here, encompassed by “qPCR”) are a critical method in environmental monitoring for a wide range of human pathogens. These assays are often selected over culture-based methods for increased sensitivity and specificity, rapid turnaround time, limited handling of biohazardous materials, and for targets that are not culturable or are difficult to culture ^1^. However, qPCR-based methods can be subject to sampling and laboratory errors leading to potential false positives and false negatives ^2–4^.

Assay signature erosion is one important source of potential false negative results, especially for viruses, which evolve more rapidly than bacterial or protozoan targets ^5^. Over time, genetic drift or natural selection can lead to fixation of mutations in regions of the target sequence where the primers or probe bind, resulting in reduced signal with a given qPCR assay ^6^. Studies examining this process found signature erosion in Ebola RT-qPCR diagnostic assays ^6^ and SARS-CoV-2 qPCR assays ^7,8^. With advances in sequencing technologies, virus genomes have been sequenced at increasing rates over time ^9^, providing an opportunity to routinely assess qPCR signature erosion using *in silico* qPCR prediction tools ^10,11^.

At the same time, PCR-based quantification of viruses from environmental samples has become widespread within wastewater-based surveillance and quantitative microbial risk assessments (QMRA). As these applications ultimately impact public health, accurate measurement is critical. We focused here on enteroviruses, a genus of non-enveloped RNA viruses (family Picornaviridae) transmitted via the fecal-oral route that are prevalent in domestic wastewater ^12^ and are a common QMRA target ^13^. Members of this diverse genus can cause an array of human illnesses including mild respiratory or diarrheal disease, hand foot and mouth disease, and acute flaccid paralysis ^14^. A variety of qPCR assays for human enteroviruses have been applied to environmental samples, ranging from EPA Method 1615, originally designed over 20 years ago ^15,16^, to more recent assays used in conjunction with sequencing ^17^. Additionally, new assays continue to be produced for clinical virology research based on sequence data ^18^, some of which may be useful for environmental surveillance. Given the importance and regular use of these assays, an assessment of signature erosion over time is warranted.

Here, we performed an *in silico* analysis of four genus-level enterovirus probe-based qPCR assays (**Table 1; Figure 1**) using tools we developed for this purpose. We examined the extent of overlap between the assays, signature erosion, comprehensiveness across enterovirus species, and the potential for false positive hits. This assessment can serve as a model for other sequence-based assay comparisons to assist in selecting and updating qPCR assays.

**Table 1.**
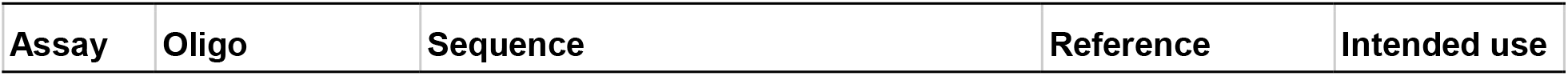
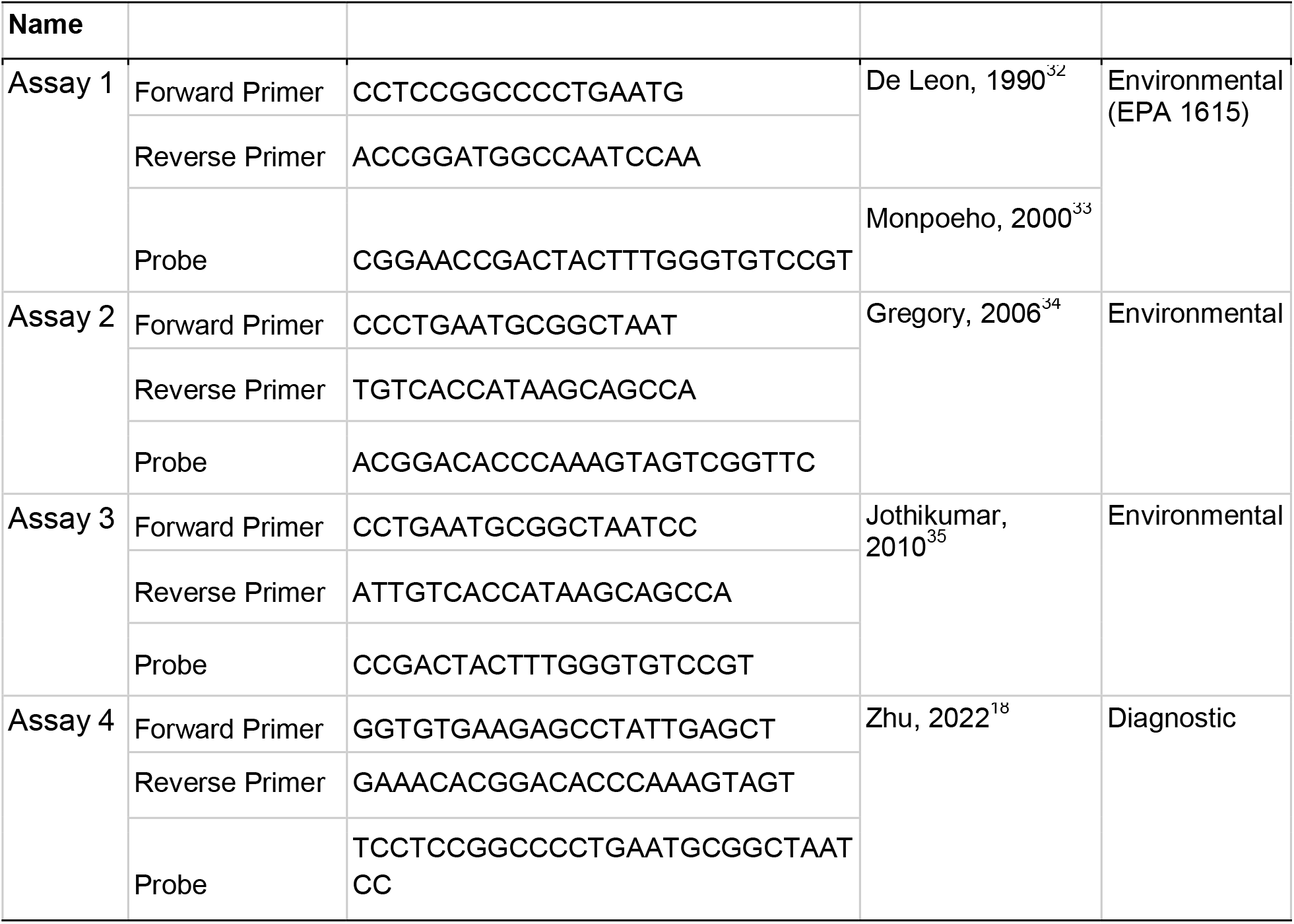
Genus-level enterovirus assays identified from literature review.

**Figure 1.**
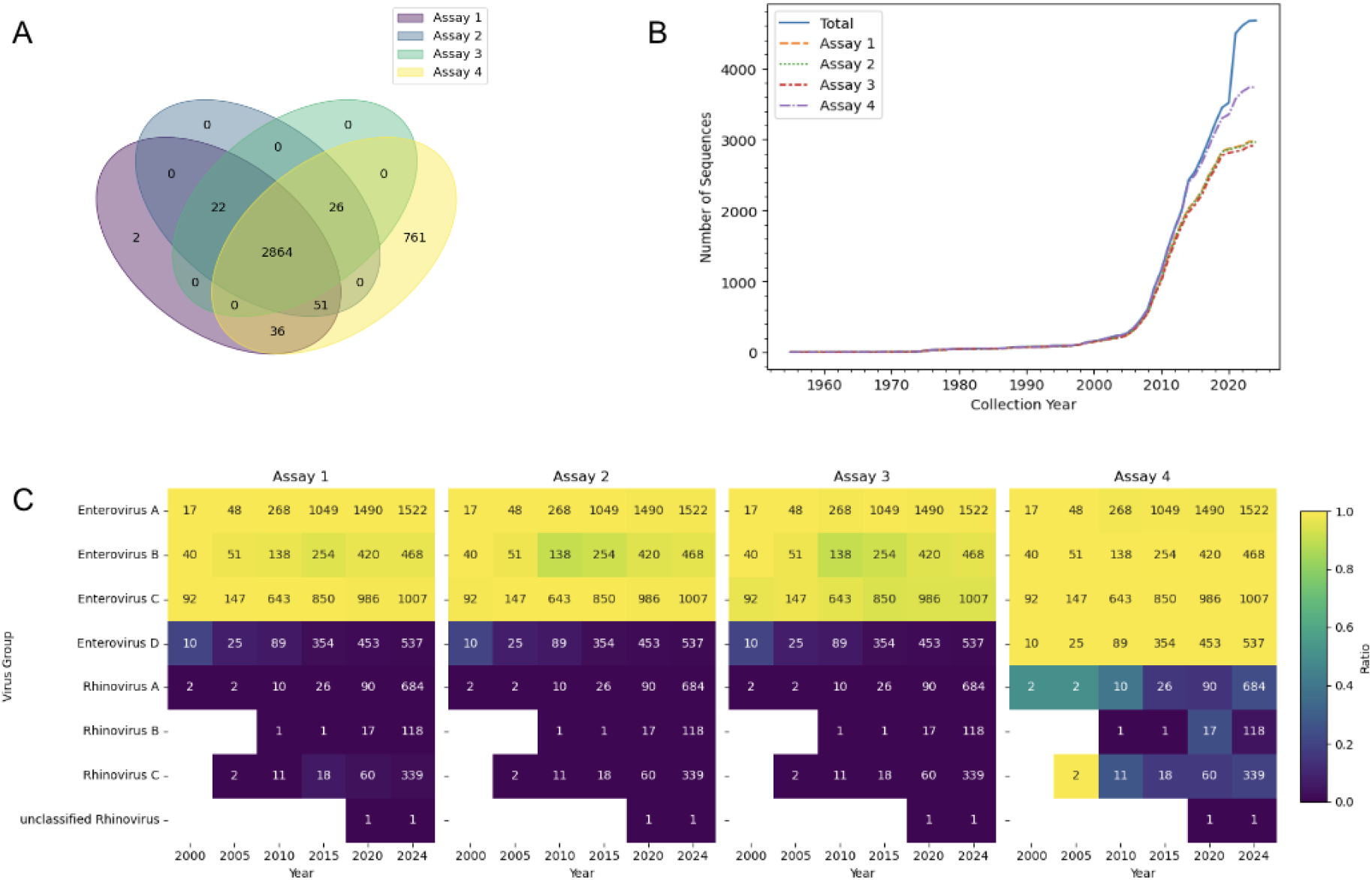
Comparison of the comprehensiveness of four pan-enterovirus assays. **A)** Venn diagram illustrating the overlap of predicted hits by each assay to the curated enterovirus database. **B)** Number of cumulative predicted hits for each assay and total genomes in the curated enterovirus database over time, based on NCBI collection date. **C)** Heatmaps showing the hit rate of each assay over time. Color indicates fraction of targets in the curated enterovirus database that were predicted to be hit by the assay. Enterovirus genomes were grouped at the species level according to their NCBI taxonomy ID. Text indicates the cumulative number of targets within each species through the year shown on the x-axis.

## Methods

### Literature Review

We searched Web of Science (Clarivate, 2024) and Google Scholar on February 29, 2024 using combinations of the keywords “qPCR”, “dPCR”, “enterovirus”, “assay”, and “assay design”. Papers were manually reviewed to identify those presenting novel assays for enterovirus (**Table 1**). A secondary search was performed to identify papers that used these assays on environmental samples (**Table S1**).

### Databases

For false negative analysis, the NCBI Datasets command line tools, *datasets* and *dataformat* (API v2, CLI v16.25.0, https://www.ncbi.nlm.nih.gov/datasets/; database accessed August 26, 2024), were used to generate a virus genome package, including metadata and genomic sequences, for all genomes from the genus “enterovirus” (TaxID 12059) in NCBI Virus.^19^ The metadata were extracted and used to filter the database for complete enterovirus genomes with the host specified as human. Genomes without a known collection date or host were excluded. Filtered sequences were clustered at 100% identity with CD-HIT (v4.8.1) to remove duplicates.^20,21^ Finally, a nucleotide BLAST database with 6149 sequences was generated.^22^

For false positive analysis, NCBI Virus (accessed August 26, 2024) was used to download accessions for all RefSeq RNA virus genomes (TaxID 2559587), excluding those that were 1000 nt or were listed as lab-passaged. Sequence data was retrieved using NCBI *datasets* and then used to generate a nucleotide BLAST database.

### *In silico* PCR prediction

We used simulate_pcr, an open-source command line tool for PCR result prediction, to analyze the four assays identified by the literature review. Inputs were qPCR assays (**Table 1**) and custom nucleotide BLAST databases (see above). For each database and each assay, simulate_pcr was run with the following parameters: minlen 50 -maxlen 1000 -mm 3 -mux 1 - num_threads 100 -max_target_seqs 1000000 -word_size 5 -evalue 100000 -3prime 3. The BLAST word size was chosen based on the length of oligos and the number of mismatches allowed based on the following reasoning: with 3 mismatches allowed per oligo and the smallest oligo length of 17 nt, the minimum exact sequence match length is 5 when the mismatches are equally spaced as far apart as possible, and any other variation gives a larger exact sequence match. The outputs from simulate_pcr were processed as described below.

### Analysis

A multiple sequence alignment of enterovirus A, B, C, and D genomes from NCBI RefSeq was performed using Geneious Prime (v2024.0.7), and primer and probe binding sites were identified in the consensus sequence ^23^. To characterize the genomes predicted to be hit and missed by each assay over time and by taxonomic group, we developed pcrvalidationtools.py, a Python 3 module that ingests data from simulate_PCR and NCBI datasets outputs (https://github.com/vannytrinh/assay_validation).

False negative analysis was complicated by the fact that many enterovirus genomes in NCBI lacked the target region of all assays. Based on the alignment, all assays of interest targeted the highly conservative 5’ untranslated region (5’UTR), which is approximately 750 nt for enteroviruses ^24^. However, genomes marked as “complete” in NCBI virus do not necessarily contain the UTR. We used the genbank records for all enterovirus genomes to determine the presence or absence of a 5’UTR: genbank records were retrieved using NCBI Entrez (through Biopython v1.81); the 5’UTR length was determined by either the length of the 5’UTR annotation (if present) or the start coordinates of the first coding sequence (CDS) annotation. Genomes with a 5’UTR <400 nt or without either 5’UTR or CDS feature annotations were excluded from the list of expected positives.

To determine the enterovirus species group for each accession, we retrieved the taxonomy record from NCBI Entrez, iterated through the genome’s taxonomic lineage, and identified the taxID directly under the enterovirus taxID.

To identify false positives, we manually reviewed predicted qPCR hits for each assay against the filtered RefSeq RNA virus database.

## Results and Discussion

Four RT-qPCR assays that broadly target the enterovirus genus were identified via our comprehensive literature review. The assays were published between 1990 to 2022 and intended for environmental and diagnostic uses (**Table 1**). Based on alignment of primers and probes to enterovirus genomes from NCBI RefSeq, all assays target the highly conserved 5’ untranslated region (5’UTR) (**Figure S1**). A secondary literature review showed that three of these assays had been applied to environmental sample types including surface water, storm runoff, municipal wastewater, wastewater primary effluent, and wastewater solids (**Table S1**).

To assess the comprehensiveness of each assay over time and across enterovirus species, we created a curated database of complete genomes from the genus “enterovirus” for which the host was specified as human. This database included sequences from seven enterovirus and rhinovirus species. We performed *in silico* PCR for all assays using this database. Of the 4676 sequences in our curated database, 2864 were predicted to be hit by all assays (**Figure 1A**). Most notably, 761 sequences were hit only by assay 4.

Next, we examined the predicted hits for each assay over time to look for evidence of signature erosion. Based on the year of collection for each genome as reported to NCBI, the number of entries in the database increased dramatically after the year 2006 (**Figure 1B**). As the database grew, the number of predicted hits also increased but then plateaued over time. Assays 1, 2, 3 exhibited similar patterns and plateaued after 2015. However, assay 4, published in 2022 using recent sequence data, captured the majority of sequences through 2020 before plateauing.

To investigate these patterns further, we determined assay coverage across four major species groups (**Figures 1C** and **S2)**. Enterovirus A was well-covered by all four assays. Coverage of enterovirus B by assays 1, 2, 3 decreased slightly, starting around 2010. Similarly, enterovirus C was well-covered, except for a slight decline in assay 3. Assay 4 was the only assay predicted to cover the enterovirus D species, however, this assay also produced more predicted hits to rhinovirus genomes than the other assays. Together, enterovirus D and rhinovirus likely accounted for the high number of non-overlapping predicted hits for assay 4 (**Figures 1A** and **S2**).

We suspect that biannual waves of enterovirus D-68 infections ^14^ and a related increase in genome submissions to NCBI contributed to the observed decrease in overall coverage by assays 1, 2 and 3 in the 2010’s (**Figures 1B** and **C**). Although these assays do not hit enterovirus D, wastewater monitoring with these assays is likely sufficient to capture large seasonal trends in enterovirus prevalence, which are largely caused by enterovirus species A, B, and C ^17^. As enterovirus D-68 is a leading cause of severe disease leading to acute flaccid paralysis, wastewater monitoring is ongoing ^25,26^ with assays for this specific strain.

Our *in silico* PCR allowed for up to three mismatches (excluding the 3’ end of primers and probes), and the occurrence of mismatches differed for each assay and each enterovirus species (**Figure S3**). Specifically, the majority of hits from assays 1, 2, and 3 had zero total mismatches between the assay (including both primers and probe) and the hit sequence (range 0 to 5). In contrast, the majority of hits with assay 4 had one or two total mismatches (range 0 to 6). There was no apparent trend in the number of mismatches across species. We note that our *in silico* PCR testing (simulate_pcr) did not consider annealing temperature, secondary structure, or primer dimers, in determining the success or failure of qPCR with each target and assay. The success of qPCR *in vitro* also depends on factors such as the qPCR reaction buffer, the enzyme, and the location and type of each mismatch ^27–29^. Future studies could compare the results from assays applied to environmental samples alongside sequencing-based methods to compare qPCR or digital PCR comprehensiveness relative to *in silico* PCR predictions.

To identify potential false positive hits with the four assays, we performed *in silico* PCR using the NCBI RefSeq virus database. Although there were no predicted hits outside of the enterovirus genus for any assay, we found eight unique predicted hits to non-human enteroviruses. Of the four assays, assay 4 had the highest number of hits, but also tended to have more mismatches to those hits (**Figure 2**). These results and the hits to rhinoviruses (**Figure 1C**) point to the need to align the goal of the analysis with the breadth of the assay: for example, assay 4 captures some non-human enteroviruses, which could be present in environmental samples. This assay may be appropriate if non-human infections are part of the surveillance goal or are expected to be at low prevalence, or if enterovirus D is a significant component of the total enterovirus in the sample. In general, care should be taken when applying human diagnostic assays to environmental samples.

**Figure 2.**
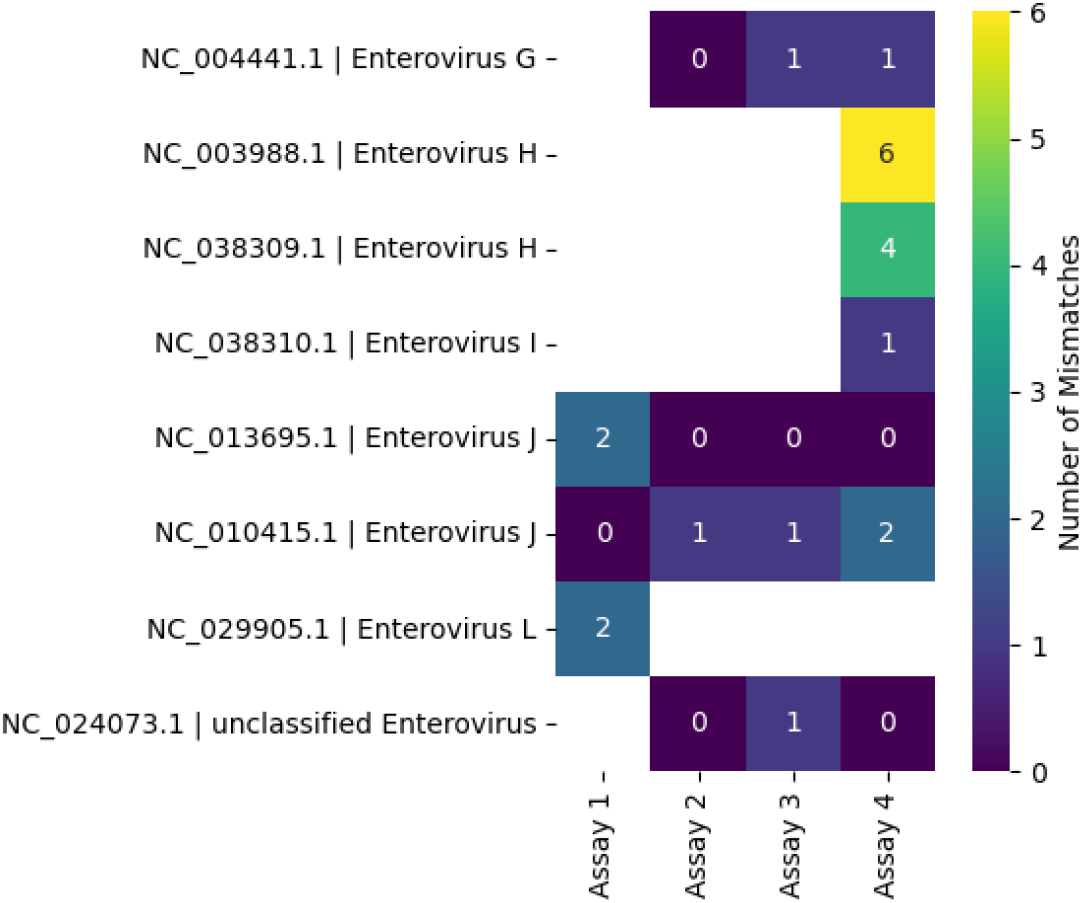
Non-human enterovirus hits from RefSeq for each assay. Color and text indicate the number of mismatches between the hit and the assay.

In addition to the limitations described above, the database affected our results. First, some enterovirus entries in the NCBI Virus database were missing information about collection date (15.7%), host (19.1%), taxonomic classification (particularly for environmental samples) and the presence or absence of the 5’ UTR. This information is left to the discretion of the submitter, and thus, careful database curation is recommended before checking qPCR assays *in silico*. Second, submissions to NCBI over time are not necessarily representative of prevalence in environmental samples and may instead reflect clinically relevant cases, one-off studies, or genomic surveillance efforts for emerging strains of concern. There may be a mismatch between clinical sequences and environmental sequences, as sequencing is conducted primarily for severe cases and may not capture the diversity of circulating viruses. Relatedly, diagnostic qPCR assays may also miss cases because the assays do not capture the circulating viruses. For example, retrospective wastewater surveillance of an outbreak of enterovirus D68 ^26^ and subsequent sequencing identified mismatches between the circulating strain and a qPCR assay that had produced false negatives in the clinic during the outbreak ^30^. As virus genomic sequencing from wastewater surveillance increases, virus databases should be systematically updated by the research and surveillance community and used to check existing and newly designed qPCR assays.

Overall, potential signature erosion and false positives in qPCR may impact the results of environmental monitoring and risk assessments. QMRA using qPCR data from out-of-date assays may over- or underestimate the risks of exposure to contaminated environmental matrices ^31^ and wastewater monitoring with an assay that is narrower than expected may miss important trends in community infections. *In silico* methods can aid users in choosing appropriate assays and in regularly checking the taxonomic coverage of those assays over time.

## Acknowledgements

Funding for this work was provided by the UCOP Laboratory Fees Research Program (award L22CR4507).

