## supplementary information for "In silico analysis reveals differential targeting of enterovirus species by commonly used PCR assays"

Tables: 1
Figures: 3

**Table S1. Environmental studies applying genus-level enterovirus assays.***

| Application | Reference | Assay |
|---|---|---|
| Urban storm water runoff and after biofilter removal | Graham, 2021[1] | 1 |
| Raw wastewater for QMRA | Pecson, 2022[2] | 1 |
| Raw sewage for water quality and QMRA models | Li, 2023[3] | 1 |
| Wastewater primary effluent to evaluate concentration methods | Brinkman, 2013[4] | 2 |
| Municipal wastewater to observe seasonal patterns | Brinkman, 2017[5] | 2 |
| Wastewater solids to compare to clinical testing | Boehm, 2024[6] | 2 |
| Seawater (*integrated cell culture qPCR protocol*) | Ming, 2011[7] | 3 |

*representative studies applying assay 1 (US EPA Method 1615) were included, but this list is not comprehensive

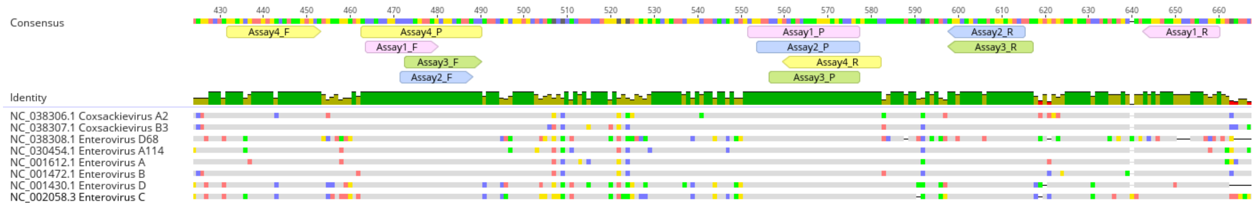


**Figure S1.** Multiple sequence alignment of the enterovirus genomes from NCBI RefSeq. qPCR assay primers ("F" and "R") and probes ("P") are annotated on the consensus sequence.

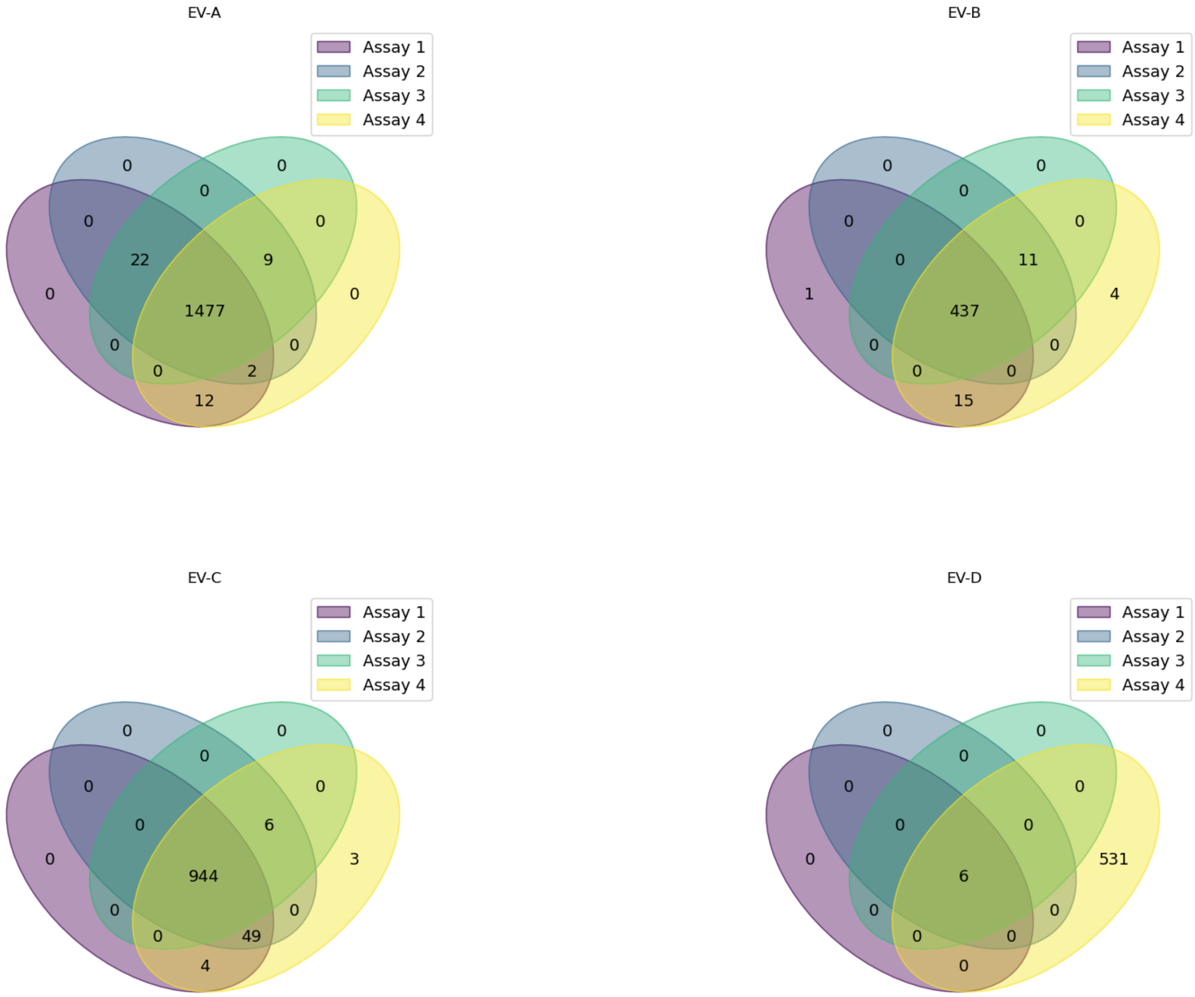


**Figure S2.** Venn diagrams illustrating the overlap of predicted hits by each assay to enterovirus species A, B, C, and D from the curated enterovirus genomes database.

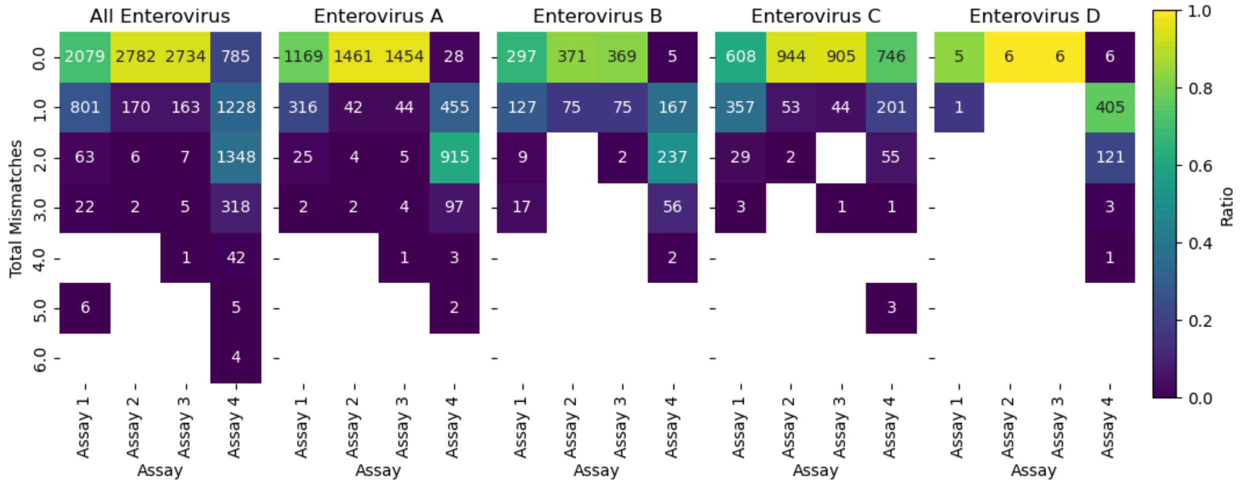

**Figure S3**. Heatmaps showing the number of total mismatches between the target and assay. Analyses are grouped by enterovirus species. Color represents the fraction of total predicted hits for each assay (x-axis) with a given number of mismatches (y-axis). Mismatch count includes all primers and probe mismatches. Text indicates the count of hits with a given number of mismatches for each assay.

(6) Boehm, A. B.; Shelden, B.; Duong, D.; Banaei, N.; White, B. J.; Wolfe, M. K. A Retrospective Longitudinal Study of Adenovirus Group F, Norovirus GI and GII, Rotavirus, and Enterovirus Nucleic Acids in Wastewater Solids at Two Wastewater Treatment Plants: Solid-Liquid Partitioning and Relation to Clinical Testing Data. *mSphere 9* (3), e00736-23. https://doi.org/10.1128/msphere.00736-23.

(7) Ming, H.-X.; Zhu, L.; Zhang, Y. Rapid Quantification of Infectious Enterovirus from Surface Water in Bohai Bay, China Using an Integrated Cell Culture-qPCR Assay. *Mar. Pollut. Bull.* **2011**, *62* (10), 2047–2054. https://doi.org/10.1016/j.marpolbul.2011.07.024.
